# Sleep Deficits and Cannabis Use Behaviors: An Analysis of Shared Genetics Using Linkage Disequilibrium Score Regression and Polygenic Risk Prediction

**DOI:** 10.1101/2020.05.02.053983

**Authors:** Evan A. Winiger, Jarrod M. Ellingson, Claire L. Morrison, Robin P. Corley, Joëlle A. Pasman, Tamara L. Wall, Christian J. Hopfer, John K. Hewitt

## Abstract

**Study Objectives:** Estimate the genetic relationship of cannabis use with sleep deficits and eveningness chronotype.

**Methods:** We used linkage disequilibrium score regression (LDSC) to analyze genetic correlations between sleep deficits and cannabis use behaviors. Secondly, we generated sleep deficit polygenic risk scores (PRSs) and estimated their ability to predict cannabis use behaviors using logistic regression. Summary statistics came from existing genome wide association studies (GWASs) of European ancestry that were focused on sleep duration, insomnia, chronotype, lifetime cannabis use, and cannabis use disorder (CUD). A target sample for PRS prediction consisted of high-risk participants and participants from twin/family community-based studies (n = 796, male = 66%; mean age = 26.81). Target data consisted of self-reported sleep (sleep duration, feeling tired, and taking naps) and cannabis use behaviors (lifetime use, number of lifetime uses, past 180-day use, age of first use, and lifetime CUD symptoms).

**Results:** Significant genetic correlation between lifetime cannabis use and eveningness chronotype (rG = 0.24, *p* < 0.01), as well as between CUD and both short sleep duration (<7 h) (rG = 0.23, *p* = 0.02) and insomnia (rG = 0.20, *p* = 0.02). Insomnia PRS predicted earlier age of first cannabis use (*β* = −0.09, *p* = 0.02) and increased lifetime CUD symptom count use (*β* = 0.07, *p* = 0.03).

**Conclusion:** Cannabis use is genetically associated with both sleep deficits and an eveningness chronotype, suggesting that there are genes that predispose individuals to both cannabis use and sleep deficits.

## Introduction

Cannabis is one of the most widely used psychoactive substances in the world ^1^ and has a well-documented but unclear relationship with sleep. Cannabis contains cannabinoids, which are the major contributors to psychoactive and medicinal effects ^2^. The brain contains cannabinoid receptors, which are affected by exogenous cannabinoids and endogenous cannabinoids (i.e. produced within the brain). Together, these receptors and endogenous cannabinoids comprise the endocannabinoid system. The two most prominent exogenous cannabinoids are Tetrahydrocannabinol (THC), the primary psychoactive cannabinoid, and Cannabidiol (CBD), which appears to have additional sedating and anxiolytic properties ^3^. Evidence suggests that an interplay between THC and CBD may underlie a nuanced relationship with sleep. For example, acute/low-dose THC and high-dose CBD may aide sleep, but low-dose CBD and long-term/high-dose THC may interfere with sleep ^4^. While cannabis is often associated with being a sleep aid ^5–10^, repeated cannabis use may lead to tolerance of its sleep-aid properties and consistent use is linked to negative sleep outcomes via habituation ^4^.

Increased frequency of cannabis use is associated with an assortment of sleep problems including prolonged latency to sleep onset ^11^, lower sleep duration ^11–16^ sleep disturbances ^17^, sleep quality problems ^17–20^, later bed times^15^, and insomnia related outcomes ^12,17,19,21–23^. These adverse effects might be specific to daily or chronic users, as a recent study found that daily cannabis users endorsed worse sleep quality and increased insomnia symptoms compared to both non-users and non-daily users, but also found that non-users and non-daily users demonstrated similar sleep scores ^17^. Thus, irregular users might not experience the adverse sleep effects experienced by heavy users. Additionally, sleep disturbances are a primary withdrawal symptom of cannabis use disorders (CUD) and are often a leading risk factor for relapse, suggesting that those with ongoing CUD might suffer from continuous sleep issues stemming from discontinued use or attempts to abstain ^4,24^. Lastly, an eveningness chronotype (a diurnal preference for a sleep-wake pattern of activity and alertness in the evening which is linked to insomnia ^25^ and sleep complaints^26 27^) is associated with increased cannabis frequency ^28–30^ and cannabis addiction ^31^.

In addition to cross-sectional association, there is evidence of early cannabis exposure and use predicting later sleep outcomes. The fetal brain is densely inhabited with CB1 receptors that spread during gestation ^32^. CB1 receptors are thought to be involved in the regulation of sleep processes since they are found in numerous regions of the brain associated with the sleepwake cycle ^33^. THC binds to CB1 receptors, and animal research implies this possibly modifies fetal cortical circuitry in the womb ^34^. Several studies have found associations of prenatal cannabis exposure with early sleep factors, such as differences in quiet time, irregular sleep, and sleep related body movements a few days after birth ^35^ and less efficient sleep and less total sleep time at three years of age ^36^.

The endocannabinoid system also plays a critical role in the development of the adolescent brain ^37^ and because the brain is changing and developing well throughout early adulthood ^38^, could be susceptible to the effects of cannabis for a large part of the lifespan. A handful of studies have found that cannabis initiation and early use predict later sleep problems such as tiredness, trouble sleeping ^22^ short sleep duration ^16,39^, and insomnia related outcomes^23^. Evidence exists for the reverse relationship as well, with premorbid insomnia ^21^ and generalized sleep problems ^39–41^ predicting later cannabis use. This effect appears strong in early development, such that early childhood sleep deficits predict cannabis use in later adolescence ^22,42–44^ and sleep factors during adolescence predict adult cannabis use ^21,45^. Lastly, endorsements of an adolescent eveningness chronotype is associated with follow-up reports of increased cannabis use controlling for baseline adolescent substance use ^40^. With evidence of both crosssectional associations and a bidirectional relationship between cannabis and sleep deficits/eveningness chronotype, there could be an underlying common liability such as shared or common genetics responsible for this association.

The concept of common genetic liability is that the same genetic influences can act on distinct or separately measured phenotypes. This can be referred to as ‘shared genetics’ or, more formally, genetic pleiotropy. That is, if phenotypes are genetically correlated, the relationship between those phenotypes can be partially explained by common genetic liability (pleiotropy, shared genetics), implying that the genetic influences on one phenotype also have an influence on another phenotype. There is increasing evidence of a genetic relationship between cannabis use and sleep deficits, which may be biologically centered on the endocannabinoid system’s involvement in the circadian sleep–wake cycle ^46–48^. Additionally, disruption of circadian genes might disturb the reward processing system, which can influence substance use ^49,50^. While research has shown evidence of common genetics between both alcohol and tobacco use and disorders with sleep outcomes using both twin studies and genomic methods (e.g., genetic correlations) ^51–56^, studies specifically focused on the genetic relationship between cannabis and sleep components remain scarce. Two twin studies have found evidence of shared genetics between cannabis use and sleep outcomes, specifically lower adult sleep duration ^16^ and adult insomnia outcomes ^23^. Additionally, several clock gene polymorphisms have been linked to risk for cannabis addiction ^57^.

Consistent with this literature, there is converging evidence that support a shared genetic liability hypothesis. Recent large genome-wide association studies (GWASs) on sleep-related and chronotype variables ^56,58–61^ have found genes and genetic pathways linked with both cannabis use or cannabinoid activity ^62–68^. Likewise, several GWASs of lifetime cannabis use and CUD disorder ^65,66,69,70^ have found genetic associations that are believed to be involved in circadian rhythm and sleep behaviors ^71–74^. These studies imply genetic pleiotropy between cannabis use and sleep deficits, but research is needed to analyze the specific role of the potential shared genetics in this relationship, specially using modern genomic methods.

As mentioned, modern GWASs have been used to identify independent genome-wide significant loci associated with various sleep traits and to estimate genome-wide single nucleotide polymorphism (SNP) heritability for several sleep-related traits such as chronotype (eveningness-morningness) (14%), sleep duration (10%), and insomnia (17%) ^56,58,75^ as well as for cannabis behaviors such as lifetime cannabis use (11%) ^66^ and CUD (4%) ^70^. These results suggest that the combined effects of common SNPs capture a considerable proportion of the heritability of various sleep behaviors and cannabis use behaviors. A polygenic risk score (PRS) is an individual measure of genetic propensity to a trait of risk that is generated by multiplying the number of risk alleles that an individual possesses at a particular SNP by the effect size from a discovery GWAS for that same SNP. By applying summary statistics from a large GWAS to a smaller genotyped target sample, PRSs can be generated to estimate if genetic risk for a trait is associated with another trait, implying shared genetics between traits. Additionally, summary statistics from GWASs can be used to analyze genetic correlations by using a technique called Linkage Disequilibrium Score Regression (LDSC) which estimates if the direction of effect of SNPs are correlated between traits. Thus, LDSC analyzes if direction of effect between the SNPs of two traits are correlated based on the whole genome, while PRS analysis uses the genetic scores assigned to individuals (derived from a GWAS) to determine if the genetic risk attributed to the PRS can predict behaviors observed in a target sample; the PRS analysis can readily control for covariates.

In this study, we first conducted genetic correlation analyses using the summary statistics of several large cannabis (lifetime cannabis use and CUD) and sleep (sleep duration, chronotype, and insomnia) GWASs with the intention of finding significant genetic correlations that would suggest that the direction of SNPs between these domains are correlated. Secondly, we generated PRSs based on summary statistics of various large sleep related GWASs (chronotype, sleep duration, and insomnia) and analyzed the ability of sleep trait PRSs to predict cannabis measures in a target sample consisting of both high-risk participants and participants from twin/family community-based studies.

## Methods

### 2.1 Participants

In order to generate accurate and bias free PRSs, the ideal analysis requires that the racial make-up of the target data reflect the original GWAS data that the PRS was derived from ^76^. With this in mind, we used an initial sample of 900 (self-identifying white) adults who participated in a third wave of data collection from either the Center on Antisocial Drug Dependence (CADD) in Boulder, CO [PI: Hewitt] or the Genetics of Antisocial Drug Dependence (GADD) cohort from Denver, CO and San Diego, CA [PI: Hopfer]. Amongst those in the sample with a family member (208 total subjects were nested within a family), we kept one member of each family at random to make our final sample 796 subjects. Removing subjects from the same family allowed us to avoid the potential convergent complications of using mixed effects models as well avoid the role of the shared environment amongst family members. Of our remaining sample, we had 644 subjects from the CADD and 152 from the GADD. Subjects from the CADD included data from the Colorado Adoption Project (n = 10; ^77^), Longitudinal Twin Study (n = 156; ^78^), Community Twin Sample (n = 315; ^78^), and the Adolescent Substance Abuse Project (n = 124; ^79^). In addition to the San Diego and Denver subsamples of the GADD, 46 subjects in this consortium were part of a separate study [PI: Hopfer, DA035804] that were integrated into the third wave of the GADD. The final sample was 66% male (n = 529) with an average age of 26.81 years (*SD* = 3.15, range = 19-37).

### 2.2 Measures

#### Cannabis measures in target sample

Cannabis measures in our target sample were self-reported via a supplement to the Composite International Diagnostic Interview Substance Abuse Module (CIDI-SAM) ^80^. Any/lifetime cannabis use was measure with a screening question regarding lifetime cannabis use (“Have you ever used cannabis?”). Subjects who endorsed any lifetime cannabis use (n = 627) were asked a series of cannabis-related questions including: age of first use (“How old were you the first time you used cannabis?”) (mean = 15.33 years, SD = 3.20), and number of lifetimes uses (“How many times in your life have you used cannabis?”). Responses for number of lifetime cannabis uses included: “1-2 times” (n = 38), “3-5 times” (n = 58), “6-9 times” (n = 28), “10-19 times” (n = 38), “20-39 times” (n = 59), and “40 or more times” (n = 416), and were coded as 1-5. Anyone who denied any lifetime cannabis use were assigned a 0 for number of lifetime cannabis use (n = 169). Previous 180-day cannabis use was measured by asking, “How many days have you used marijuana in the past six months (180 Days)?” (mean = 34.52 years, SD = 67.62). Past 180-day cannabis use was categorized as 0 days (n = 562), 1-100 days (n = 189), and more than 100 days (n = 149).

We included several measures of DSM-IV cannabis use disorders ^81^ taken from the CIDI-SAM in order to generate a measure that consisted of the sum of the number of both lifetime cannabis dependence and abuse symptoms endorsed (mean = 1.52, SD = 2.51, range = 0-11), with the goal of generating a variable conceptually similar to the unidimensional symptom count of CUD in DSM-5 ^82^. Research has determined that the cannabis abuse and dependence criteria of DSM-IV might not distinguish between two separate disorders or constructs ^83^ and that a unidimensional symptom count classifies CUD more appropriately ^84^. Studies with DSM-IV cannabis dependence and abuse symptom data have utilized this summation technique in order to make a symptom count measure that is comparable to CUD reflecting a single disorder ^84,85^.

#### Sleep measures in target data

Sleep measures in our target sample were assessed using the Jessor Health Questionnaire ^86^. Sleep duration was assessed using two questions which asked, “How many hours of sleep do you typically get on a weekend?” and “How many hours of sleep do you typically get on a weekday?” with responses being “5 hours or less,” “6 hours,” “7 hours,” “8 hours,” “9 hours,” “10 hours,” or “11 hours or more.” Our measures of short sleep duration were coded to match the GWAS we generated our PRS from ^56^. Short sleep duration <7 hours was coded as a 1 (n = 233 and n = 162) and 7–8□hours sleep duration was coded as 0 (n = 500 and n = 415) for both weekday and weekend sleep respectively. Those who reported 9 or more hours were assigned an NA in order to match the coding of the sleep duration GWAS. Subjects also were asked: “How often do you feel tired or sleepy when you get up in the morning?”, “How often do you feel tired or low on energy in day?”, and “How often do you take a nap during the daytime?” with possible responses for these three questions being “almost never,” “once a week or so,” “2 or 3 times a week,” “nearly every day,” and “would rather not answer;” coded as 1-4 and NA.

#### Generating polygenic risk scores

After removing duplicate SNPs, quality control performed in PLINK ^87^ included pruning variants based on missingness (>5%), minor allele frequency (<1%), and Hardy-Weinberg equilibrium (*p*-value <.001) followed by pruning variants based on linkage disequilibrium (wherever r^2^ exceeds .20 within a 50 kb window). There were 1,089,148 SNPs available for generating PRSs after applying QC. Effect sizes from the discovery GWAS of sleep traits ^56,58,75^ were multiplied by the number of affected alleles at each individual SNP in our target sample in order to generate a unique PRS for each participant. That is, each participant in our data was assigned a PRS for each sleep trait, by applying GWAS effect sizes to their genomic data. Each PRS comprised of all SNPs that passed quality control steps (all *p* < 1) and thus explained the highest variance in the sleep phenotypes in our target sample. This is consistent with work suggesting that complex traits display a high amount of polygenicity and that using only genome-wide significant SNPs may exclude many SNPs with small but meaningful additive effects ^88^; thus including all possible SNPs captures the highest amount of variance possible for a given trait ^89,90^.

#### Summary statistics from sleep GWASs for PRS analysis and genetic correlations via LDSC

Sleep trait PRS effect sizes were generated from the summary statistics of several large sleep-related GWASs (all performed by the same research team) that utilized data of European ancestry from the UK biobank ^56,58,75^. The sleep-related traits included self-report chronotype ^58^, short sleep duration (<7 hours) ^56^, and self-reported insomnia symptoms ^75^. The original chronotype GWAS utilized morningness, which was a binary measure of being a morning person or not (if participants endorsed any morningness measures as opposed to any eveningness measure) with morning people coded as 1 and evening people coded as 0. For the purpose of this study and the emphasis on sleep deficits, we reverse coded this measure and interpreted the results as eveningness chronotype (127,622 cases and 120,478 controls). Short sleep was defined as <7□hours relative to 7–8□hours sleep duration (106,192 cases and 305,742 controls). Severe Insomnia (as defined by the GWAS) was classified using a self-report question: “Do you have trouble falling asleep at night or do you wake up in the middle of the night?”, with participants being dichotomized into controls (“never/rarely”, n = 108,357) and frequent insomnia symptoms (“usually”, n = 129,270) and with those reporting “sometimes” excluded. These same summary statistics were used for the genetic correlation analysis.

#### Summary statistics from cannabis GWASs for genetic correlations via LDSC

Summary statistics from several large scale GWASs of cannabis use behaviors were used for LDSC. Summary statistics for lifetime cannabis use were generated from self-report measures of whether a participant had ever used cannabis during their lifetime from a sample comprised of both the UK biobank and the International Cannabis Consortium (n□=□162,082 ^66^). Summary statistics for CUD were derived via the International Statistical Classification of Diseases and Related Health Problems, 10th revision diagnosis of CUD ^91^ reflecting a problematic and persistent use of cannabis and were derived from the deCODE cohort based in Iceland (5,501 cases and 301,041 controls) ^70^.

#### Covariates

We included age ^92^, sex ^93,94^, depression ^95,96^, and current alcohol and tobacco use ^97,98^ as covariates in all regression models. Current alcohol and tobacco use were measured using the number of days that tobacco (mean = 35.09 SD = 43.25) and alcohol (mean = 60.05, SD = 80.59) were used in the past 180-days. Past 180-day tobacco use was categorized as 0 days (n = 123), 1-10 days (n = 196), 11-40 days (n = 246), and more than 40 days (n = 231). Past 180-day alcohol use was categorized as 0 days (n = 386), 1-100 days (n = 163) and more than 100 days (n = 247). Depression symptoms were assessed using the Center for Epidemiological Studies-Depression (CES-D) scale ^99^ with the caveat of the sleep disturbance question being removed due to its direct overlap with our sleep PRS measure (mean = 27.66, SD = 8.96). All regression models also included the first 10 ancestral principal components (PCs 1-10) generated in PLINK as covariates to account for population stratification as is common in Genetic and PRS analysis ^100,101^.

### 2.3 Statistical Analysis

#### Genetic correlations via linkage disequilibrium score regression

We used Linkage Disequilibrium Score Regression (LDSC) ^102^ to calculate genetic correlations between traits. Summary statistics were filtered by INFO > .90 and MAF > .01. Strand ambiguous SNPs, SNPs with duplicated rs numbers, and multi-allelic variants that are not SNPs were all removed. SNPs with low Ns (as determined by the LDSC program) were also removed when sample sizes were available. Alleles were merged with the Hap Map 3 ^103^ reference panel, with the major histone complex removed. The LD scores and beta weights used were pre-computed from 1000 Genomes European GWAS data included in the LDSC download.

LDSC is a computationally efficient method that regresses Chi-square statistics from GWASs on LD scores of the trait of interest ^104^. LD scores per SNP are the sum of the variance explained by LD with other SNPs ^105^. Genetic correlations were calculated using overlapping SNPs from filtered summary statistic files. Genetic correlations also account for population stratification and are not confounded by overlapping samples.

#### Polygenic risk regression analysis

All regression analyses were conducted in R version 3.5.1 ^106^. Logistic regression models were used to test the association between sleep trait PRSs and cannabis use behaviors including the previously mentioned covariates of age, sex, depression symptoms, past 180-day alcohol and tobacco use, and ancestral principal components (PCs 1-10). In terms of the steps of our analyses, we first used LDSC to estimate potential genetic correlations between sleep and cannabis GWAS summary statistics. Second, we ran phenotypic regression models between our sleep measures and cannabis behaviors in our target data to establish associations of sleep deficits and cannabis behaviors amongst our target sample. Third, we ran regression models between our sleep PRSs and sleep traits in our discovery sample in order to confirm that the sleep PRSs predicted sleep constructs in our target data. Lastly, we ran regression models to see how sleep PRSs predict cannabis behaviors. For all series of regressions models involving PRSs we ran two sets of models 1) with just age, sex, and PCs 1-10 as covariates and 2) with the prior covariates and the addition of current depression symptoms and past 180-day alcohol and tobacco use. We utilized this two-model approach in order to 1) look at the associations between sleep PRSs and cannabis factors controlling for both basic covariates and more complex covariates and 2) determine if the effects seen in our final models (including all covariates) were driven by the more maladaptive covariates (depression and past 180-day substance use).

## Results

### Genetic correlations using LDSC

We first used LDSC to look at potential genetic correlations between cannabis and sleep traits using the largest GWAS to date for each trait. Table 1 displays the LDSC analysis between sleep traits and cannabis use traits. We found significant positive genetic correlations between any lifetime cannabis use and eveningness chronotype (rG = 0.24, *p* < 0.01). We also found significant genetic correlations between CUD and both short sleep duration (rG = 0.23, *p* = 0.02) and insomnia (rG = 0.20, *p* = 0.02), as well as a marginally significant genetic correlation between CUD and eveningness chronotype (rG = 0.16, p = 0.06). Figure 1 displays the genetic correlations between cannabis and sleep phenotypes using LDSC (Error bars are 95% confidence intervals).

**Table 1.** Genetic correlations between sleep and cannabis phenotypes using large scale GWAS.

|  | Lifetime Cannabis Use | Cannabis Use Disorder |
| --- | --- | --- |
| Short Sleep Duration (<7 h) | -0.05 [0.03] | **0.23\* [0.10]** |
| Eveningness Chronotype | **0.24\* [0.03]** | 0.16# [0.09] |
| Insomnia | 0.01 [0.03] | **0.20\* [0.09]** |

*Note.* Genetic correlations between cannabis and sleep phenotypes were calculated using LD Score Regression. Standard errors are in brackets.
\*indicates $p < 0.05$ , # $p = 0.06$

### Sleep traits predicting cannabis in target data

Table 2 displays regression outputs of our target data sleep traits predicting cannabis use behaviors controlling for sex, age, depression, and past 180-day alcohol and tobacco use. Short sleep duration on the weekday significantly predicted earlier age of first cannabis use (*β* = −0.06, *p* < 0.01). Short sleep duration on the weekend significantly predicted increased number of lifetime cannabis uses (*β* = 0.23, *p* < 0.01), increased past 180-day cannabis us (*β* = 0.24, *p* < 0.01), and earlier age of first cannabis use (*β* = −0.07, *p* < 0.01), and was trending for lifetime cannabis use (*β* = 0.51, *p* = 0.09). How often one feels tired or sleepy in the morning significantly predicted increased lifetime CUD symptom count (*β* = 0.09, *p* = 0.01). How often one felt tired or low on energy during the day significantly predicted early age of first cannabis use (*β* = −0.09, *p* = 0.02), but was trending in predicting increased lifetime CUD symptom count (*β* = 0.07, *p* = 0.09).

**Table 2.** Regression betas of sleep traits predicting cannabis use behaviors controlling for sex, age, depression, and past 180-day substance use.

| **Sleep Trait** | Lifetime Cannabis Use | Number of Lifetime Uses | Age of First Cannabis Use | Past 180-day Cannabis Use | CUD Symptom Count |
| --- | --- | --- | --- | --- | --- |
| How often do you feel tired or sleepy when you get up in the morning? | 0.11 | 0.12 | -.03 | -0.04 | **0.09\*** |
| How often do you feel tired or just low in energy during the day? | 0.11 | 0.12 | **-0.09\*** | -0.01 | 0.07# |
| How often do you take a nap during the daytime? | 0.12 | 0.16 | -0.08# | 0.08# | 0.09# |
| Short sleep duration on the weekday (<7 h) | 0.01 | 0.04 | **-0.06\*\*** | 0.04 | 0.04 |
| Short sleep duration on the weekend (<7 h) | 0.51# | **0.23\*\*** | **-0.07\*\*** | **0.24\*\*** | 0.04 |

### Sleep PRSs predicting sleep traits

Table 3 displays the regression betas between the sleep PRSs and our sleep variables in our target data controlling for age, sex, and PCs 1-10. We found evidence of significant associations between the PRS for short sleep duration and numerous sleep factors including short weekday sleep duration (*β* = 0.32, *p* < 0.01), short weekend sleep duration (*β* = 0.23, *p* = 0.04), and frequency of taking naps during the day (*β* = 0.08, *p* = < 0.01). Eveningness chronotype PRS was significantly associated with how often one feels tired or low in energy during the day (*β* = 0.07, *p* = 0.03). While the Insomnia PRS trended in predicting short sleep duration on the weekday (*β* = 0.15 *p* = 0.07), we failed to find significant associations between the insomnia PRS with any of our sleep measures. Table 4 displays regression betas for the sleep PRSs predicting sleep outcomes/traits in our target data in our full models controlling for age, sex, PCs 1-10, current depression, and past 180-day alcohol and tobacco use. Short sleep PRS significantly predicted short sleep on the weekday (*β* = 0.34, *p* < 0.01), short sleep on the weekend, (*β* = 0.27, *p* = 0.02), and how often one takes naps during the day (*β* = 0.08, *p* < 0.01). Eveningness chronotype significantly predicted how often one feels tired or low in energy during the day (*β* = 0.08, *p* = 0.01). The insomnia PRS did not significantly predict any sleep variables in our target data but was trending for predicting short sleep duration on the weekday (*β* = 0.16, *p* = 0.07).

**Table 3.**
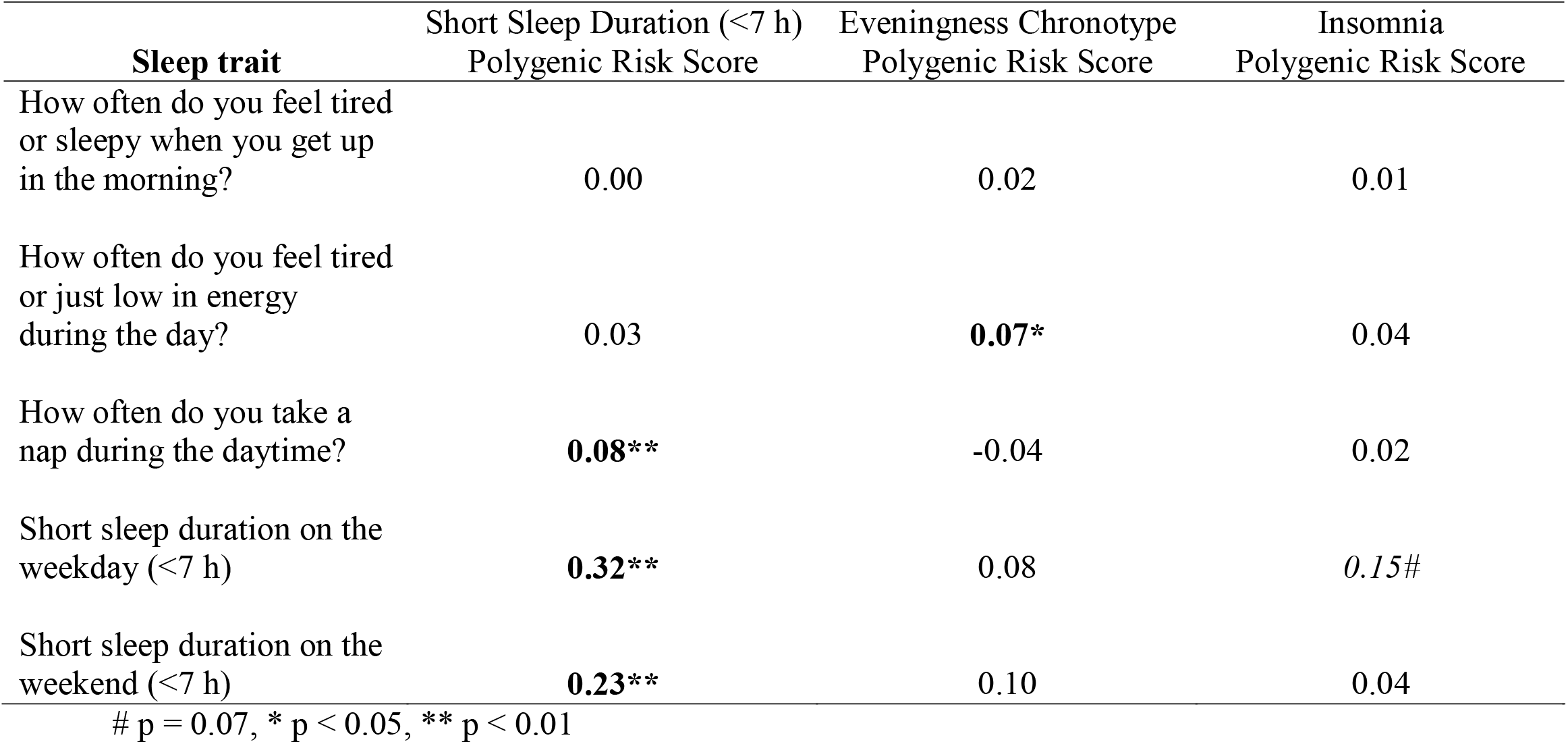
Regression betas for sleep polygenic risk scores (*p* < 1) predicting sleep factors controlling for age, sex, and ancestral principal components (PCs 1-10).

**Table 4.** Regression betas for sleep polygenic risk scores (*p* < 1) predicting sleep factors controlling for sex, age, depression, past 180-day substance use, and ancestral principal components (PCs 1-10).

| **Sleep trait** | Short Sleep Duration (<7 h)<br>Polygenic Risk Score | Eveningness Chronotype<br>Polygenic Risk Score | Insomnia<br>Polygenic Risk Score |
| --- | --- | --- | --- |
| How often do you feel tired or sleepy when you get up in the morning? | 0.00 | 0.03 | 0.02 |
| How often do you feel tired or just low in energy during the day? | 0.04 | **0.08\*** | 0.05 |
| How often do you take a nap during the daytime? | **0.08\*\*** | 0.04 | 0.02 |
| Short sleep on the weekday (<7 h) | **0.34\*\*** | 0.10 | *0.16#* |
| Short sleep on the weekend (<7 h) | **0.27\*** | 0.09 | 0.05 |

### Sleep PRSs predicting cannabis traits

Table 5 displays the regression betas between the sleep PRSs and our sleep variables in our target data controlling for age, sex, and PCs 1-10. The insomnia PRS significantly predicted earlier age of first cannabis use (*β* = −0.11, *p* = 0.01) and increased lifetime CUD symptom count (*β* = 0.08, *p* = 0.02). Short sleep duration PRS was trending for predicting increased past 180-day cannabis use (*β* = 0.07, *p* = 0.07), as was the eveningness chronotype PRS for predicting lifetime cannabis use (*β* = 0.15, *p* = 0.09) and increased lifetime CUD symptom count (*β* = 0.06, *p* = 0.09). Table 6 displays the regression betas for the sleep PRSs predicting cannabis use measures in our target data controlling for age, sex, PCS 1-10, current depression, and past 180-day alcohol and tobacco use. The insomnia PRS significantly predicted earlier age of first cannabis use (*β* = −0.09, *p =* 0.02) and increased lifetime CUD symptom count (*β* = 0.07, *p* = 0.03). Both the short sleep duration and eveningness chronotype PRSs did not significantly predict any of our cannabis measures.

**Table 5.** Regression betas for sleep polygenic risk scores (*p* < 1) predicting cannabis behaviors controlling for sex, age, and ancestral principal components (PCs 1-10).

| **Sleep trait** | Short Sleep Duration (<7 h)<br>Polygenic Risk Score | Eveningness Chronotype<br>Polygenic Risk Score | Insomnia<br>Polygenic Risk Score |
| --- | --- | --- | --- |
| Lifetime Cannabis Use | 0.10 | *0.15#* | 0.09 |
| Number of Lifetime Cannabis Uses | 0.02 | 0.05 | 0.04 |
| Age of First Cannabis Use | -0.06 | 0.00 | **-0.11\*\*** |
| Past 180-day Cannabis Use | *0.07#* | 0.03 | 0.01 |
| CUD Symptom Count | 0.02 | *0.06#* | **0.08\*** |

**Table 6.** Regression betas for sleep polygenic risk scores (*p* < 1) predicting cannabis behaviors controlling for sex, age, depression, past 180-day substance use, and ancestral principal components (PCs 1-10).

| **Cannabis Trait** | Short Sleep<br>Duration (<7 h)<br>Polygenic Risk<br>Scores | Eveningness<br>Chronotype<br>Polygenic Risk<br>Score | Insomnia Polygenic<br>Risk Score |
| --- | --- | --- | --- |
| Lifetime Cannabis Use | 0.10 | 0.16 | 0.15 |
| Number of Lifetime Uses | -0.01 | 0.05 | 0.04 |
| Age of first Cannabis Use | -0.04 | 0.00 | **-0.09\*** |
| Past 180-day Cannabis Use | 0.06 | 0.03 | 0.01 |
| CUD Symptom Count | 0.01 | 0.05 | **0.07\*** |

\* $p < 0.05$

## Discussion

We set out to examine the potential shared genetic liability of sleep deficits and cannabis use behaviors using multiple genomic methods. We found significant genetic correlations between lifetime cannabis use and eveningness chronotype as well as significant positive genetic correlation between CUD and both short sleep duration and insomnia. Additionally, we found that an insomnia PRS generated from a large scale sleep GWAS predicted earlier age of first cannabis use and increased number of lifetime CUD symptoms controlling for sex, age, PCs 1-10, current depression, and past 180-day alcohol and tobacco, suggesting that the genetic risk attributed to insomnia can predict several cannabis use behaviors. This study presents the first genomic based evidence (using PRS and LDSC) of shared genetic influence for cannabis use behaviors with both sleep traits and chronotype. The direction of these results implies a shared genetic relationship between increased cannabis use behaviors and both sleep deficits and an eveningness chronotype

Our LD score derived genetic correlations are the first such report of a genetic relationship between sleep deficits and cannabis use behaviors, and are similar to prior studies that have shown genetic associations of sleep deficits and other substance use behaviors such as alcohol and tobacco ^54–56^. These results and analyses imply that there is a positive correlation between the effect of SNPs between sleep deficits (short sleep duration, insomnia, and an eveningness chronotype) and lifetime cannabis use/CUD, such that the genetic influences on cannabis use also have an influence on sleep deficits or vice versa. While a prior twin study found a genetic correlation between eveningness chronotype and both increased alcohol quantity and binge drinking ^51^, to our knowledge this is the first report of chronotype being genetically correlated with cannabis use. Additionally, our PRS analysis findings present the first report of genetic risk for a sleep behavior predicting a substance use behavior, suggesting that the genetic risk for insomnia is associated with cannabis use behaviors such earlier age of first cannabis use and increased number of lifetime CUD symptoms.

Our sleep duration PRS was validated on phenotypes of sleep behaviors in our target sample. Specifically, our short sleep duration PRS predicted short sleep duration on the weekday/weekend and how often one takes naps during the daytime. Additionally, the eveningness chronotype PRS predicted how often one feels tired or low in energy during the day. Unfortunately, our target sample did not have a direct measure of insomnia and our regression analysis failed to find any significant relationships between the insomnia PRS and the sleep outcomes in our target data (although there was a trending association between the insomnia PRS and short sleep duration on the weekday). Similarly, the only chronotype-like measure was restricted to a question regarding how often one feels tired or sleepy in the morning, which differed from the morning person/night person question of the chronotype GWAS. The insomnia GWAS phenotype was defined as severe insomnia, excluding those who had insomnia symptoms “sometimes” and only including those with “usual/always” insomnia symptoms. This exclusion could have led the PRS to predict only extreme cases and this could have influenced the potential variance explained in our regression analysis regarding the sleep outcomes.

It is worth noting that we included cannabis measures from differential time points with the goal of not only analyzing the genetic relationship between sleep and various cannabis behaviors across life, but also to look at the phenotypic associations of early cannabis use and later sleep in our target data. We found that several measures of recent sleep characteristics were associated with earlier cannabis use measures in our target data; for example, short sleep duration on the weekend was associated with earlier age of first cannabis use as well as increased number of lifetime uses, and both short sleep duration on the weekday and how often one feels tired or low in energy during the day were associated with earlier age of first cannabis use. Additionally, we found that increased lifetime CUD symptom count was associated with feeling tired or sleepy in the morning. These results support prior findings of potentially earlier or preceding cannabis use behaviors being associated with later sleep factors. Lastly, our measure of past 180-day cannabis use was significantly associated with short sleep duration on the weekend, implying phenotypic associations of recent cannabis use with recent sleep measures.

While some of the results from our LDSC analysis align with our PRS analysis (e.g., findings of shared genetics between insomnia and CUD) several of our findings were not replicated between the two analyses. For instance, we found a genetic correlation between short sleep duration and CUD, yet our PRS for short sleep duration did not predict any cannabis behaviors. Additionally, we found a significant genetic association between eveningness chronotype and lifetime cannabis use, yet the eveningnness chronotype PRS did not predict any cannabis behaviors. Reasons for the lack of convergent results could include population differences between the GWAS and target data in terms of both environmental and genetic differences of the samples. Different locations of the samples will have different environmental influences that can influence phenotype expression and there are both racial and regional differences in terms of common and rare variants, minor allele frequencies, and linkage disequilibrium that can influence results and the variance explained ^107^. Lastly, differences in the methodological aspects of the analyses (PRS vs LDSC) could be responsible. While both analyses focused on the effects across all available SNPs, LDSC looks at the overall direction of effect of all SNPS and PRS looks whether the genetic risk for a certain trait predicts a phenotype. Overall, our results imply shared genetics between cannabis use and sleep deficits, and the differences seen in the results may be due to population and methodological differences between these analyses.

Our findings complement a small collection of research focused on the genetics of cannabis use and sleep behaviors. Two prior studies by our group have used the classical twin design to show that shared genetics played a role in the etiology of the relationship between early cannabis use and shorter adult sleep duration, insomnia, and insomnia with short sleep ^16,23^ and a recent study found clock gene polymorphisms that were significant risk factors for cannabis addiction ^57^. One possible explanation for this genetic relationship could be that disturbances of circadian rhythm genes might interrupt the reward processing system, which could influence substance use ^49,50^. Another supported explanation could be that the endocannabinoid system is involved in the circadian sleep-wake cycle, such that endocannabinoids influence sleep behaviors and their levels can vary with the time of day and other circadian related factors ^46–48,108^. Along these lines, several of the genes and genetic pathways found to be significant in sleep related-variable GWAS ^56,58–61^ have been associated with cannabis use and cannabinoid activity ^62–65,67,68,109–112^. Likewise, GWAS for lifetime cannabis use and CUD ^65,66,69,70^ have found genes that have been linked to sleep behaviors and circadian rhythm ^71–74,113^.

This study demonstrates a genetic relationship between both sleep factors and chronotype with cannabis use behaviors, implying shared genetic liability between these domains, specifically common genetics between short sleep duration, insomnia, and eveningness chronotype with increased cannabis use behaviors. Future studies should use more novel genetic methods to examine the exact mechanisms for this genetic relationship such as gene set enrichment pathway analysis ^114^. There are also mechanisms outside of genetics that could be responsible for the associations between these traits and future research should use methods that can make causal inferences like Mendelian randomization ^115^ and epigenome-wide association studies ^116^ to study the relationship between cannabis use and sleep deficits.

### Limitation

There are several limitations to this study, which point to important lines of future research. First, our cannabis, sleep, and covariate variables in our target sample were self-report and could be prone to response bias or report error. Second, while both the short sleep duration and eveningness chronotype PRSs were significantly associated with sleep duration outcomes and other sleep behaviors, the insomnia PRS was not significantly associated with any of our target data sleep measures (although trending for weekday sleep duration). The cohort study lacked a valid insomnia or chronotype measure to validate the usefulness of the respective evening chronotype and insomnia PRS measures. Inclusion of such measures would have ideally been associated with these PRS measures. Third, population differences in environmental factors between target and base data can influence the variance predicted in the models. Our GWAS data for our PRS was gathered from the UK in a cohort known for being older, predominantly female, overtly healthy, and mostly white ^117^. Our target sample was also of European ancestry but was younger, mostly male, and a combination of community-based and high-risk subjects. Fourth, genetic differences due to the regional make-up our samples could influence the variance. While our GWAS data for our PRSs was from European ancestry/predominantly white and our target cohort was made up of only subjects who self-identified as white, there could still be genetic differences between the samples that could have influenced the predictive ability of the PRSs to explain the variance of the outcomes. Fifth, we reported the effects of all SNPs (*p* < 1) in our results, and while using this threshold method captures the additive effect of additional SNPs often removed by the stringent threshold of genome wide significance ^88–90^, it is also susceptible to false positives or noise. Still, studies have shown that the whole-genome approach of using all SNPs captures more signal than it does noise and that this method can outperform PRSs generated from using only top hits ^118,119^. Sixth, while LDSC is robust to population stratification/relatedness there are limitations to consider such as biases in the estimates due to rare copy variants and capturing genetic variation tagged only by common SNPs ^102^. Lastly, several of our phenotypic and genetic based results regarding the relationship between cannabis use and sleep deficits were trending in significance and it is possible that similar studies with considerably larger samples would yield clearer results.

### Summary

Our findings are consistent with the theory that both sleep deficits (such as short sleep duration and insomnia) and eveningness chronotype share genetic liability with cannabis use behaviors, and that this genetic relationship contributes to the associations between sleep and cannabis. These results extend the current body of research focused on the relationship of sleep and cannabis behaviors to include the first instance of genomic evidence (LDSC and PRS prediction) as well as the first evidence of a genetic relationship between eveningness chronotype and cannabis use behaviors. Future studies should consider novel genomic methods to examine potential genes as well as specific genetic causal pathways for these relationships.

---

**Figure 1**. Genetic correlations between cannabis and sleep phenotypes. Genetic correlations were calculated with LDSC. Error bars are 95% confidence intervals.

